# Checkpoint kinase 1 is essential for establishing fetal haematopoiesis and hematopoietic stem and progenitor cell survival

**DOI:** 10.1101/413310

**Authors:** Fabian Schuler, Sehar Afreen, Claudia Manzl, Georg Häcker, Miriam Erlacher, Andreas Villunger

**Affiliations:** Division of Developmental Immunology, Biocenter, Medical University of Innsbruck, Innsbruck, Austria; Division of Pediatric Hematology and Oncology, Department of Pediatrics and Adolescent Medicine, Faculty of Medicine, University of Freiburg, Freiburg, Germany; Faculty of Biology, University of Freiburg, Freiburg, Germany; Department of Pathology, Medical University of Innsbruck, Innsbruck, Austria; Institute of Medical Microbiology and Hygiene, University Medical Center Freiburg, Freiburg, Germany; German Cancer Consortium (DKTK), Freiburg, Germany and German Cancer Research Center (DKFZ), Heidelberg, Germany

**Keywords:** CHK1, BCL2, apoptosis, DNA-damage, hematopoietic stem cells

## Abstract

Checkpoint kinase 1 is critical for S-phase fidelity and preventing premature mitotic entry in the presence of DNA damage. Tumour cells have developed a strong dependence on CHK1 for survival and hence this kinase has developed into a popular drug-target. *Chk1*-deficiency in mice results in blastocyst death due to G2/M checkpoint-failure showing that it is an essential gene and may be difficult to target therapeutically without side-effects. Here, we show that chemical inhibition of CHK1 kills murine hematopoietic stem and progenitor cells (HSPCs) as well as human CD34^+^ HSPCs by the induction of BCL2-regulated but p53-independent apoptosis. Moreover, *Chk1* is essential for stem cell survival and definite hematopoiesis in the mouse embryo. Remarkably though, cell death inhibition in hematopoietic stem cells (HSC) cannot restore blood cell formation *in utero* as *Chk1* loss causes severe DNA damage that ultimately prevents HSC expansion. Our findings establish a previously unrecognized role for CHK1 in establishing hematopoiesis; they also suggest adverse effects of therapeutic CHK1-inhibtion, particularly under conditions forcing stem cells out of dormancy, such as chemotherapy-induced myelosuppression.

## INTRODUCTION

The hematopoietic system originates from one specialized cell type, the hematopoietic stem cell (HSC). Remarkably, HSCs are not the first blood cells that can be detected in mammals. Indeed, at around embryonic day 7.5 (E7.5) in mice the first blood cells can be detected in so-called blood-islands, which are found in extra-embryonic tissue like the placenta, the vitelline duct and the yolk sac (1-4). These blood cells, primitive erythrocytes and macrophages, arise from primitive HSC (pHSC) that have limited properties in colony formation assays and are unable to reconstitute the adult blood system in bone-marrow transplantation experiments (5, 6). In contrast, definite HSC (dHSC) are capable of establishing the entirety of all blood cells of an organism and have self-renewal capacity for life. Both, pHSC and dHSC develop from a common ancestor, an Etv2^+^ Flk1^+^ mesodermal cell with endothelial and hematopoietic potential (6-8). RUNX1 and GATA transcription factors dictate their fate. GATA1^+^ RUNX1^+^ cells define pHSC that provide the embryo with primitive erythrocytes and macrophages until birth when blood formation starts to rely entirely on dHSC (9). Cells that down-regulate RUNX1 commit as “angioblasts” to the endothelial lineage. Together with GATA1^-^ RUNX1^+^ cells, angioblasts contribute to the formation of the hemogenic endothelium at E8.0. Lineage tracing experiments have shown that GATA1^-^ RUNX1^+^ cells commit to the hematopoietic lineage and are therefore the first dHSC detectable in the dorsal aorta of the aorta-gonad-mesonephros (AGM) at E9.5-10.5 (10). From there, dHSC migrate to, expand and initiate the first wave of definite hematopoiesis in the fetal liver between E11.5-E14.5 (11) and are dependent on GATA2 for survival (12). Shortly before birth, dHSC migrate further to the bone marrow to reside there in specific niches protected from environmental threats but responsive to cues that define the need to replenish the hematopoietic system (13, 14).

Long-term HSC (LT-HSC) in the adult have self-renewal capacity but reside in a quiescent state for most of their lifetime (15). LT-HSCs divide asymmetrically into one daughter with short-term (ST) reconstitution potential, differentiating into a multi-potent progenitor (MPP) cell and the other one remaining a *bona fide* stem cell (16). MPPs then commit to the myeloid, lymphoid or erythroid/megakaryocyte lineage. These transient amplifying progenitors with limited lineage potential provide the organism with the blood cells needed. To fulfil this task, the cell cycle of LT-HSCs and their immediate progeny is tightly regulated intrinsically but also responds to signals from the bone marrow micro-environment. Macrophages, for example, are thought to influence Nestin^+^ mesenchymal stem cells (MSCs) to stop secreting stem cell factor (SCF), angiopoietin-1, CXCL12 and vascular cell adhesion molecule (VCAM) 1, to induce cell cycle entry in HSCs (17). Furthermore, regulatory T-cells, Schwann cells and osteoblasts were reported to locally inhibit HSC expansion (18-20). Intrinsically, HSC-quiescence is promoted by the polycomb-protein BMI1 and the p53 tumor suppressor (15, 21).

The serine/threonine kinase checkpoint kinase 1 (CHK1) is a critical cell cycle regulator that controls normal proliferation and is activated in response to DNA-damage, thereby also controlling p53 function (22, 23). Especially upon single-stranded DNA breaks, ataxia-telangiectasia and Rad3-related protein (ATR) phosphorylates CHK1, leading to its activation and stabilisation (24). On the one hand, active CHK1 arrests the cell cycle via inhibition of CDC25 phosphatases that are essential for the activity of Cyclin/CDK-complexes. CHK1 phosphorylated CDC25A is marked for ubiquitination and therefore protesomal degradation leading to a drop in CDK2-activity and subsequent G1/S-arrest (25, 26). On the other hand, CDC25C is retained in the cytoplasm by 14-3-3 proteins when phosphorylated by CHK1 upon DNA damage, restraining CDK1-activity leading to a G2/M-arrest (27). Moreover, CHK1 promotes the activity of MYT1 and WEE1 kinases that both inhibit CDK1 by phosphorylation blocking transition from G2 to M-phase (28, 29). Under these conditions, CHK1 can stabilize p53 by direct phosphorylation to tighten cell cycle arrest (30, 31). In the absence of p53, however, cells become highly dependent on CHK1 for cell cycle control, arrest and repair of DNA damage (24, 26), generating a vulnerability that is currently explored as a means to treat cancers with CHK1 inhibitors (23, 32).

*Chk1* deletion in mice was shown to be embryonic lethal around E5.5 due to G2/M checkpoint failure. Blastocysts lacking *Chk1* exhibit massive DNA damage and cell death that could not be overcome by co-deletion of *p53* (33, 34). All of this supports the essential function of *Chk1* in cell cycle regulation and the DNA damage response to avoid mutational spread and genomic instability. Of note, a certain percentage of *Chk1*^*+/-*^ mice was reported to develop anaemia with age, suggesting critical dose-dependent roles in erythropoiesis (35) while conditional deletion of *Chk1* in B and T cells was shown to arrest their development at early proliferative stages due to accumulation of DNA damage and increased cell death (36, 37). This suggests that blood cancer treatment with chemical inhibitors targeting CHK1 may cause collateral damage within the healthy hematopoietic system, at least in cycling lymphoid or erythroid progenitors, yet the role of *Chk1* in early hematopoiesis and stem cell dynamics remains unexplored.

It was reported that *Chk1* mRNA is expressed at significant levels in HSC (35) despite the fact that HSC remain quiescent for the majority of their lifetime. Given the fact that HSC accumulate DNA damage when exiting dormancy (38, 39), e.g. under pathological conditions such as substantial blood loss or in response to infection (40-42), as well as during natural aging (43, 44), it appears appropriate that HSCs arm themselves with CHK1 to immediately deal with the dangers of a sudden proliferative challenge to avoid mutational spread. Yet, another study found that *Chk1* mRNA levels are low in HSC but increase during proliferation-coupled self-renewal or differentiation, along with other DNA damage response and repair genes (43). Consistent with a direct link to proliferation, *Chk1* mRNA was also found to be higher in fetal liver *vs.* bone-marrow derived HSC from young or aged mice (43). Notably here, HSC from old mice suffer from enhanced replication stress upon mobilization that triggers substantial CHK1 activation (44). Together this suggests critical roles for CHK1 in stem cell dynamics and prompted us to elucidate its role in the development and survival of the hematopoietic stem and progenitor cells. Using a conditional allele for *Chk1* in combination with a CRE-deleter strain allowing recombination of this allele within the hematopoietic system, we were able to highlight the importance of *Chk1* for the establishment of hematopoiesis in the embryo and the survival of HSPCs from mice and man.

## RESULTS

### CHK1 inhibition limits the survival of hematopoietic progenitors by inducing mitochondrial apoptosis

To explore the consequences of CHK1 inhibition on pluripotent hematopoietic precursors we first assessed the response of Hoxb8-immortalized FLT3-dependent progenitor cells (Hoxb8-FL cells). These cells can be generated from murine bone marrow and are multi-potent progenitor (MPP)-like cells that can be differentiated *in vitro* into myeloid and lymphoid lineages (45). Treatment of Hoxb8-FL cells with two specific CHK1 inhibitors (CHK1i), PF-477736 (PF) or CHIR-124 (CHIR) resulted in time and dose-dependent cell death, as assessed by propidium iodide (PI) staining of DNA-content by flow cytometric analysis (Figure 1A, Figure S1). Hoxb8-FL cells generated from *Vav-BCL2*-transgenic mice or mice lacking the key-apoptotic effectors *Bax* and *Bak* were protected from cell death indicating initiation of caspase-dependent mitochondrial apoptosis after CHK1i treatment (Figure 1A). The rapid cell death seen in wild type cells was accompanied by γH2AX accumulation, p53-stabilization, and PARP1 cleavage, as detected in western analyses (Figure 1B). As γH2AX phosphorylation is also induced during apoptosis due to CAD-mediated cleavage of DNA between nucleosomes (46), we wondered if this may be a secondary consequence of caspase-activation. Consistently, upon CHK1 inhibition, Hoxb8-FL cells lacking BAX/BAK also stabilized p53, showed significantly lower levels of γH2AX phosphorylation but no signs of PARP1 cleavage or cell death (Figure 1B). This suggests that CHK1i treatment causes DNA damage, triggering a p53-response that precedes BCL2-regulated mitochondrial cell death involving caspase activation. Remarkably though, Hoxb8-FL cells derived from the bone marrow of *p53*^*-/-*^ mice were still highly susceptible to cell death indicating a minor contribution of p53 target genes to apoptosis upon CHK1 inhibition (Figure 1A).

**Figure 1:**
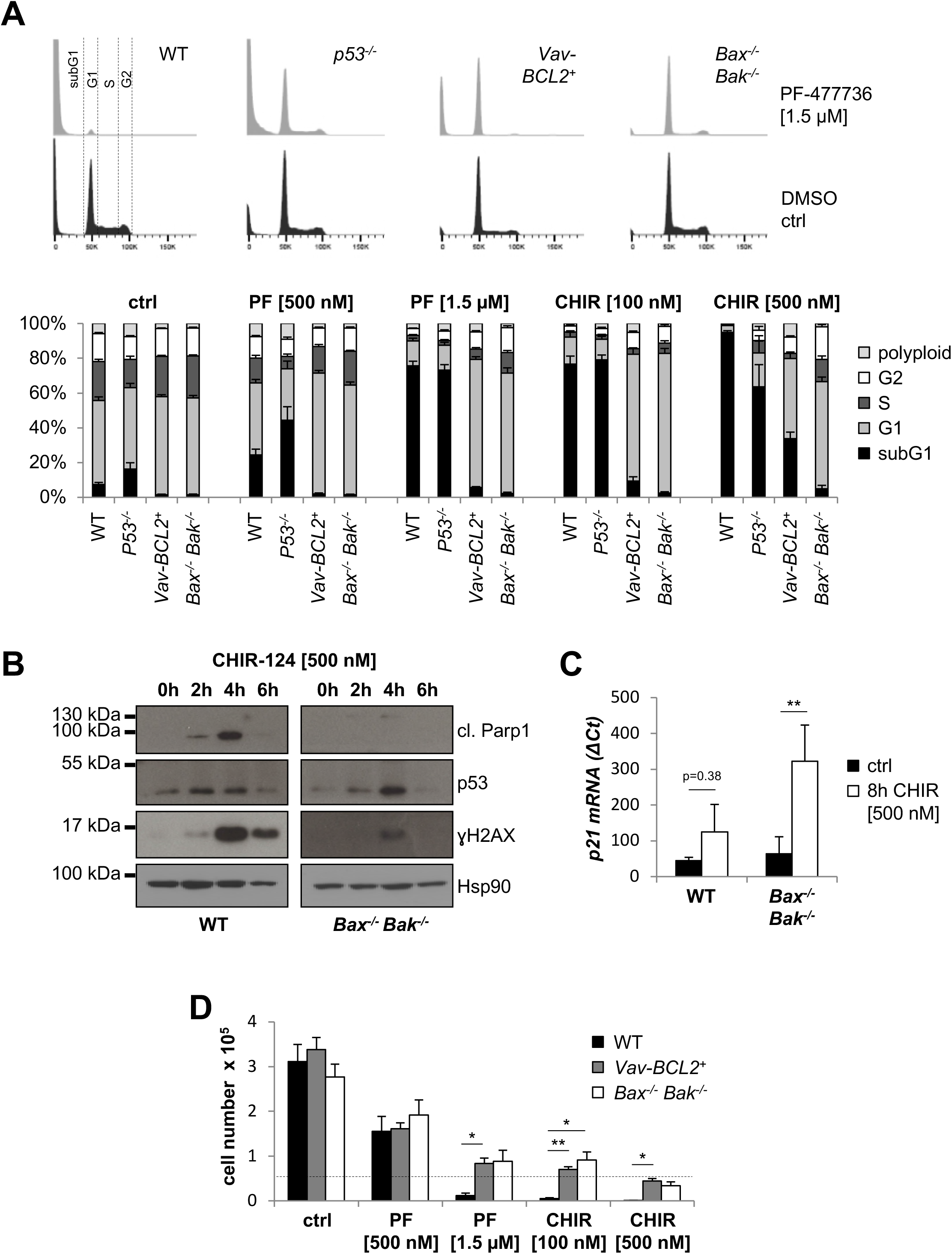
CHK1 is an essential survival factor for Hoxb8-FL cells. **(A)** Hoxb8-FL cells of the indicated genotypes were treated for 24h with graded doses of the CHK1-inhibitors PF-477736 (PF) or CHIR-124 (CHIR). Representative histograms of CHK1i-treated (grey) or DMSO-treated (black) cells analysed for cell cycle distribution and cell death using Nicoletti staining and flow cytometry are shown (Propidium Iodide, PI). Bars represent means ±S.E.M. (n=3 biological replicates per genotype). **(B)** Wild-type (WT) and *Bax*^*-/-*^ *Bak*^*-/-*^ Hoxb8-FL cells were treated for 2, 4 and 6h with 500 nM CHIR and processed for **w**estern blotting using the indicated antibodies. **(C)** WT, and *Bax*^*-/-*^*Bak*^*-/-*^ Hoxb8-FL cells were treated for 8h with 500 nM CHIR. Shown here are the relative increases in *p21* mRNA levels compared to *Hprt* in response to CHK1-inhibition, as assessed by real-time RT-qPCR.**(D)** 50.000 Hoxb8-FL cells of the indicated genotypes were seeded to assess population doublings in response to CHK1-inhibition. Cells were counted 48h post seeding. Bars represent means ± S.E.M. (n=3 biological replicates per genotype). Bars represent means ± S.D. of two experiments. Asterisks indicate significant differences: *p < 0.05, **p < 0.01, ***p < 0.001 using unpaired Student’s t-test.

Interestingly, we also noted an altered cell cycle distribution in cells that failed to undergo cell death upon CHK1-inhibition. The majority of BCL2-overexpressing or BAX/BAK-deficient cells accumulated in the G1 fraction with a near complete loss of S-phase cells, indicating cell cycle arrest at the G1/S boundary when CHK1 function was inhibited and cell death blocked at the same time (Figure 1A,B). Consistently, BAX/BAK-deficient cells showed a significant induction of *p21* mRNA in response to CHK1 inhibition (Figure 1C). To corroborate this finding, we also performed population-doubling experiments in the presence or absence of CHK1-inhibitor. Indeed, cells overexpressing BCL2 or cells lacking BAX/BAK failed to proliferate when treated and showed near-constant cell numbers over time (Figure 1D).

Together these findings underscore the importance of CHK1 for hematopoietic progenitor cell proliferation and survival by preventing BAX/BAK-mediated apoptotic cell death. Apoptosis seems initiated in response to DNA-damage upon CHK1-inhibition but does not require p53. When cell death is blocked, these cells show signs of DNA damage and arrest the cell cycle at the G1/S boundary.

### CHK1-inhibition kills primary mouse and human HSPCs via BCL2-regulated apoptosis

To get a first impression how CHK1 inhibition affects primary HSPCs, we isolated Lin^-^ Sca1^+^ cKit^+^ (LSK) cells, containing HSCs and MPPs, from the fetal liver of WT or *Vav-BCL2* transgenic embryos or the bone marrow of adult mice of the same genotypes. Sorted LSK cells from either source showed only modest cell death when treated with CHK1i but the cell death seen was blocked when BCL2 was overexpressed (Figure 2A). This suggests that, compared to Hoxb8-FL cells, their reduced proliferative capacity impacts on sensitivity to CHK1i. Consistently, when we treated total fetal liver (FL) or total bone marrow cell cultures that both contain a substantial number of lineage-committed cycling hematopoietic progenitors with CHK1i, we observed a dose-dependent response, driving a significant portion of cells into apoptosis. Again, cell death was significantly reduced when BCL2 was overexpressed (Figure 2B,C). This suggests that actively cycling hematopoietic progenitors are more vulnerable, compared to LSK cells, suggesting that cell cycle rates define susceptibility to CHK1i.

**Figure 2:**
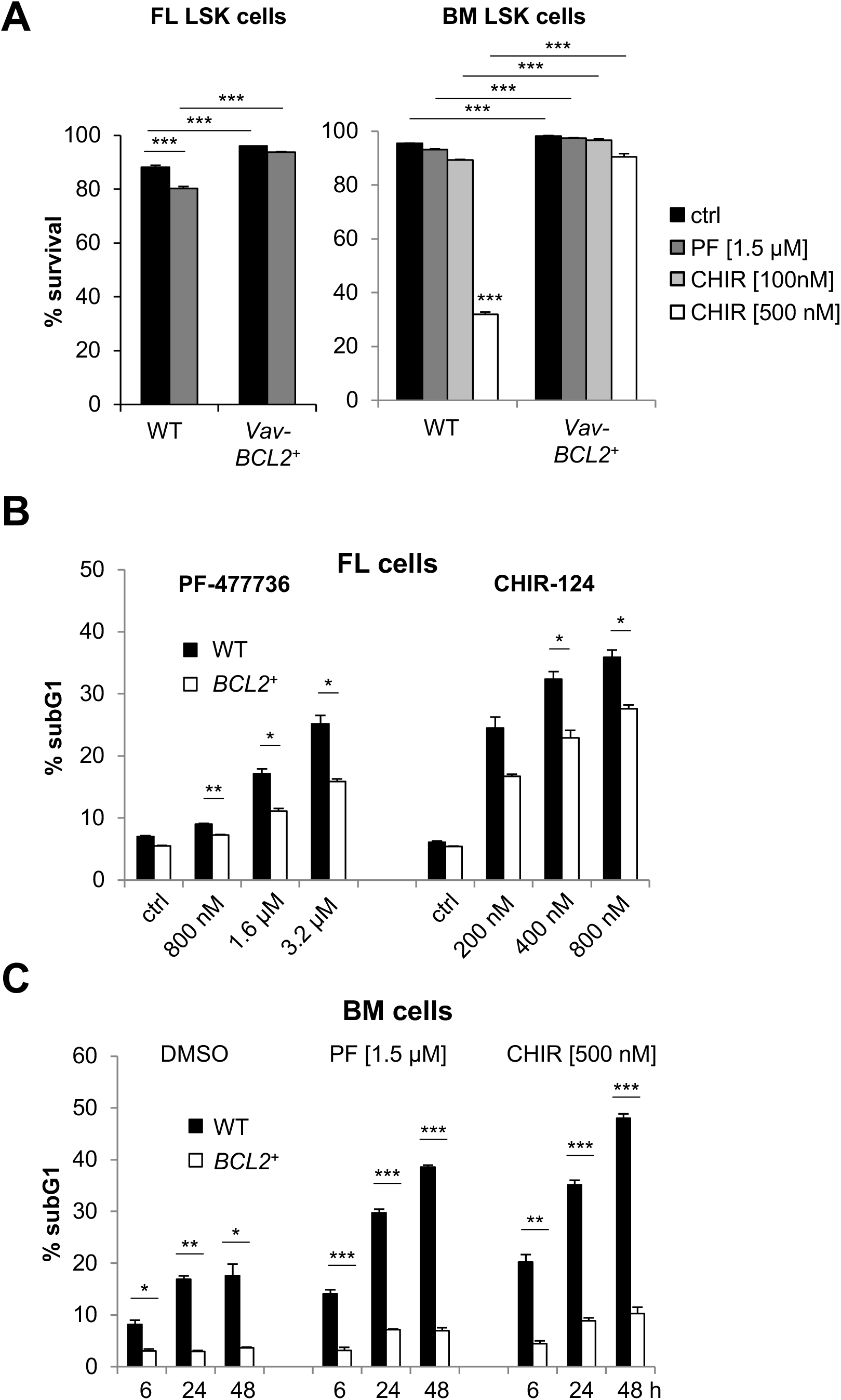
CHK1-inhibition kills primary murine hematopoietic stem and progenitor (HSPC) cells. **(A)** LSK (Lin^-^ Sca1^+^ cKit^+^) cells of the indicated genotypes were isolated by cell sorting from the fetal liver at embryonic day E13.5 or the bone marrow of adult mice and treated for 48h with PF-477736 [1.5 µM]. Survival was assessed using AnnexinV-staining and flow cytometry. Bars represent means ± S.E.M. from n=6 wild type and n=3 Vav-BCL2 embryos (left) and n=3 adults/genotype (right). WT and Vav*-BCL2* derived E14.5 total fetal liver cells **(B)** or total bone marrow cells **(C)** were treated for 72h with different doses of CHK1i. Survival was assessed using Nicoletti staining and flow cytometry. Bars represent means ± S.E.M. (n=3 biological replicates per genotype). Asterisks indicate significant differences: *p < 0.05, **p < 0.01, ***p < 0.001 using unpaired Student’s t-test.

To test whether these observations also hold true for human cells, we purified CD34^+^ HSPCs from the cord-blood that reportedly express low level CHK1 (47). Indeed, a significant fraction of CD34^+^ cells died when treated with increasing doses of CHK1i in the presence of proliferation-inducing cytokines (Figure S1A). This cell death was potently blocked by the pan-caspase inhibitor QVD pointing towards mitochondrial cell death (Figure 3A). In line with our findings in mouse cells, cell death was reduced when BCL2 was introduced into CD34^+^ cells by viral gene transfer (Figure 3B) and led to an enrichment of GFP^+^ cells, marking viral transduction, upon treatment (Figure 3C, Figure S1B). Moreover, human CD34^+^ HSPCs lost their potential to form colonies in methyl-cellulose when CHK1 was inhibited chemically (Figure S1C). In line with our experiments using Hoxb8-FL cells, BCL2 overexpression failed to rescue colony formation upon CHK1i inhibition (Figure 3D).

**Figure 3:**
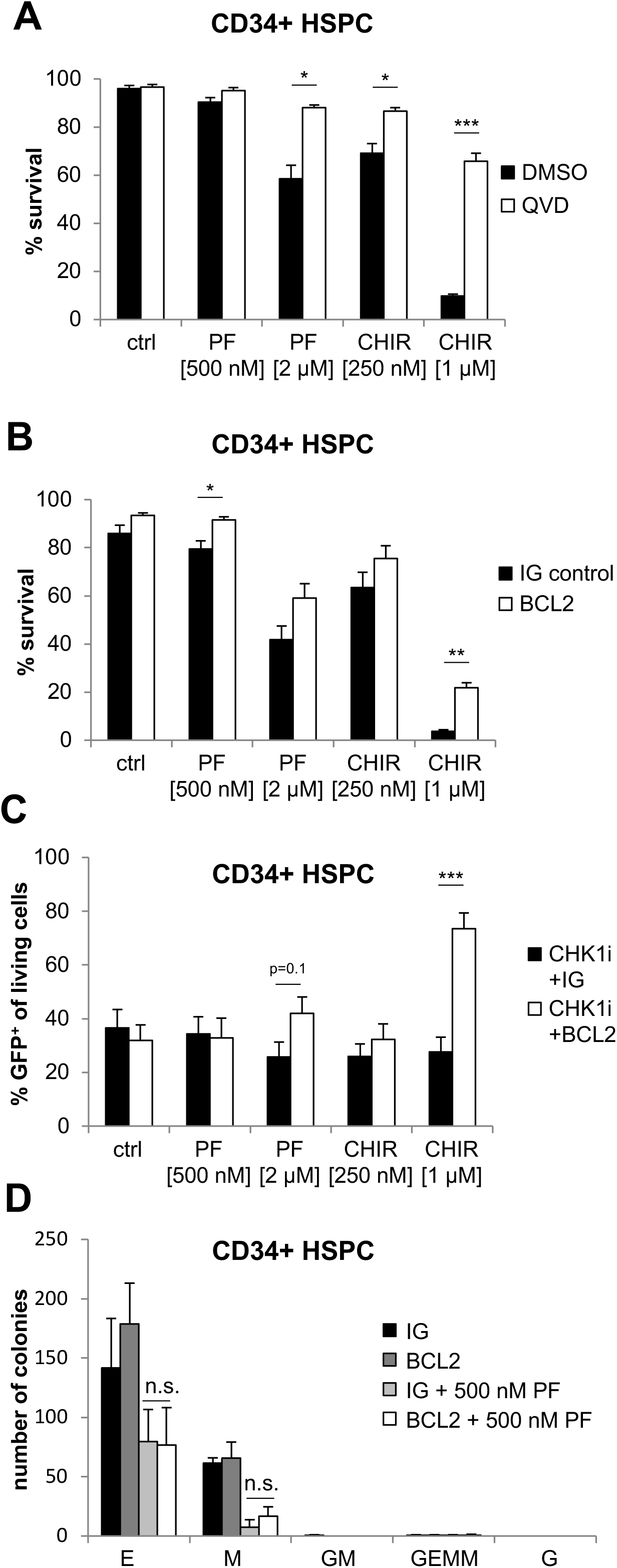
CHK1-inhibition kills human cord-blood derived CD34^+^ hematopoietic stem and progenitor (HSPC) cells in a BCL2 regulated manner. **(A)** MACS-purified CD34^+^ human cord-blood derived HSPC were treated with PF-477736 or CHIR-124 ± the caspase-inhibitor QVD (50 µM). Cell death was assessed using AnnexinV/7-AAD staining and flow cytometry (n=4 independent experiments). **(B)** CD34^+^ cells were transduced with empty vector or a vector encoding BCL2, along with IRES-GFP with an MOI of 10 for two consecutive days, followed by CHK1i application for 48 hours. Survival was assessed using Nicoletti staining and flow cytometry. Bars represent means ± S.E.M. (n=4 independent experiments). (**C)** Percentage of living GFP^+^ virus-transduced human HSPCs, cultured in the absence or presence of CHK1i for 48 hours. **(D)** Colony formation potential of human CD34^+^ HSPC transduced with empty control or BCL2 encoding virus. Colonies were counted 10 days post seeding of 1000 CD34^+^ cells per plate/sample. E: Erythroid, GM: granulocyte + monocyte, M: monocyte, G: granulocyte, GEMM: granulocyte, erythroid, monocyte/macrophage and megakaryocyte. N=4 donors. ± S.E.M. Asterisks indicate significant differences: *p < 0.05, **p < 0.01, ***p < 0.001 using unpaired Student’s t-test.

Together, this suggests that BCL2-regulated and caspase-dependent apoptosis is initiated also in CD34^+^ human HSPCs treated with CHK1-inhibitor expanded *ex vivo* and that blocking cell death is insufficient to restore their clonal outgrowth, most likely because of cell cycle arrest, or, less likely, induction of non-apoptotic cell death.

### CHK1 is essential for establishing hematopoiesis in the fetal liver

Our data indicate that CHK1 might be critical also for the development of a functional hematopoietic system. To explore this genetically, we crossed mice harboring a conditional allele of *Chk1* with a CRE-deleter strain specific for the hematopoietic system (48). Deletion of *Chk1* using *Vav*-driven CRE expression never resulted in viable mice at the time of weaning but caused hematopoietic failure and the death of *Chk1*^*fl/-*^*Vav-Cre* embryos *in utero* (Figure 4A,B). The lack of red blood cells in the fetal liver of *Chk1*^*fl/-*^*Vav-Cre* embryos, seen already at E12.5 (not shown), was easily visible histologically (Figure 4C,D) and pointed towards impaired erythropoiesis. Indeed, flow cytometric analysis revealed a decreased percentage of pro-erythroblasts (CD71^+^ cKit^+^) as well as erythroid progenitor (CD71^+^ CD45^low^) cells in the fetal liver of *Chk1*^*fl/-*^*Vav-Cre* embryos and the erythrocytes present in these embryos appeared to be of primitive origin (49), based on the lack of cKit expression on their cell surface (Figure 4E,F). Furthermore, we could observe a reduction of pro-erythrocytes (ProE, CD71^+^ Ter119^-^) and a trend of a higher number of mature Ter119^+^ erythrocytes, EryC, combined with a significant reduction of EryB cells in the fetal liver of *Chk1*^*fl/-*^*Vav-Cre* embryos.

**Figure 4:**
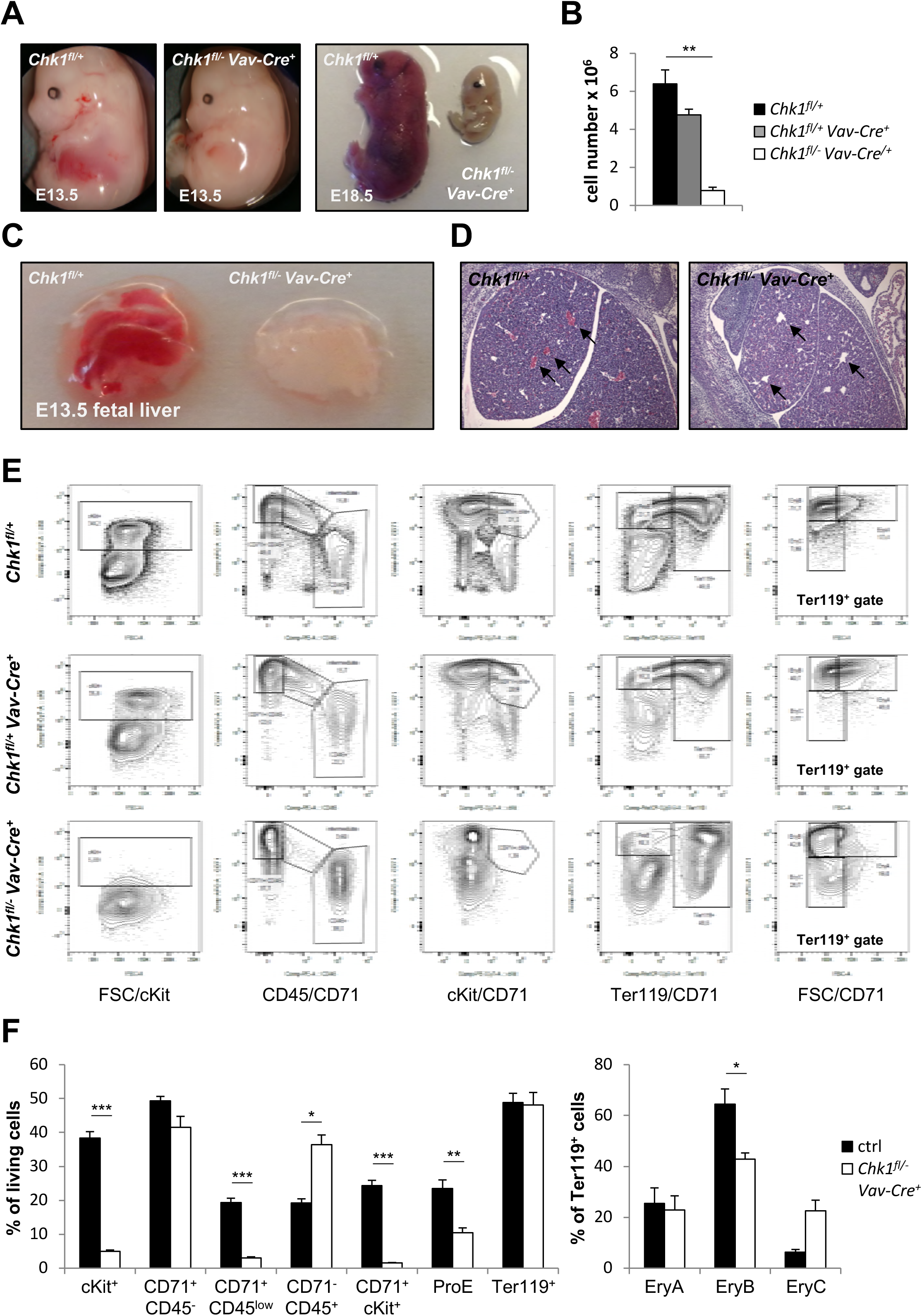
Deletion of *Chk1* in HSC prevents fetal hematopoiesis. **(A)** Representative pictures of E13.5 and E18.5 embryos of the indicated genotypes. **(B)** Quantification of fetal liver cell number at E13.5. N=4 for *Chk1*^*fl/+*^, N=3 for *Chk1*^*fl/+*^ *Vav-Cre*^*+*^ and N=4 for *Chk1*^*fl/-*^ *Vav-Cre*^*+*^. **(C)** Representative pictures of isolated fetal livers at E13.5. **(D)** Representative H&E-staining of fetal liver sections from E13.5 embryos. Black arrows indicate lack of erythroid cells in the blood vessels of *Chk1*^*fl/-*^ *Vav-Cre*^*+*^ fetal livers. **(E)** Single cell suspensions of fetal livers from *Chk1*^*fl/+*^, *Chk1*^*fl/+*^ *Vav-Cre*^*+*^ and *Chk1*^*fl/-*^ *Vav-Cre*^*+*^ embryos were stained with antibodies recognizing ckit, CD71, Ter119 or CD45 to assess primitive and definite erythropoiesis by flow cytometry. **(F)** Quantification of analyses shown in (E). Bars represent means ± S.E.M. ctrl N=5 (N=4 *Chk1*^*fl/+*^ and N=1 *Chk1*^*fl/+*^ *Vav-Cre*^*+*^), N=3 for *Chk1*^*fl/-*^ *Vav-Cre*^*+*^. Asterisks indicate significant differences: *p < 0.05, **p < 0.01, ***p < 0.001 using unpaired Student’s t-test.

Of note, we also failed to detect the typical LSK cell population enriched for HSPCs in the fetal liver. Within the Lin^-^ fraction of cells we found instead Sca1^+^ cells with a near 10-fold reduction of cKit expression (LSK^low^ cells), a phenomenon not seen in littermate controls (Figure 5A,B). Remarkably, the percentage of LSK cells was actually increased about 5-fold leading to comparable absolute cell numbers as found in littermate controls. In contrast, the fraction of Lin^-^ LK cells, containing MPPs and more committed progenitors, was significantly reduced, both in relative and absolute terms (Figure 5A,B). This suggested that LK cells cannot survive or expand properly in the absence of *Chk1*.

**Figure 5:**
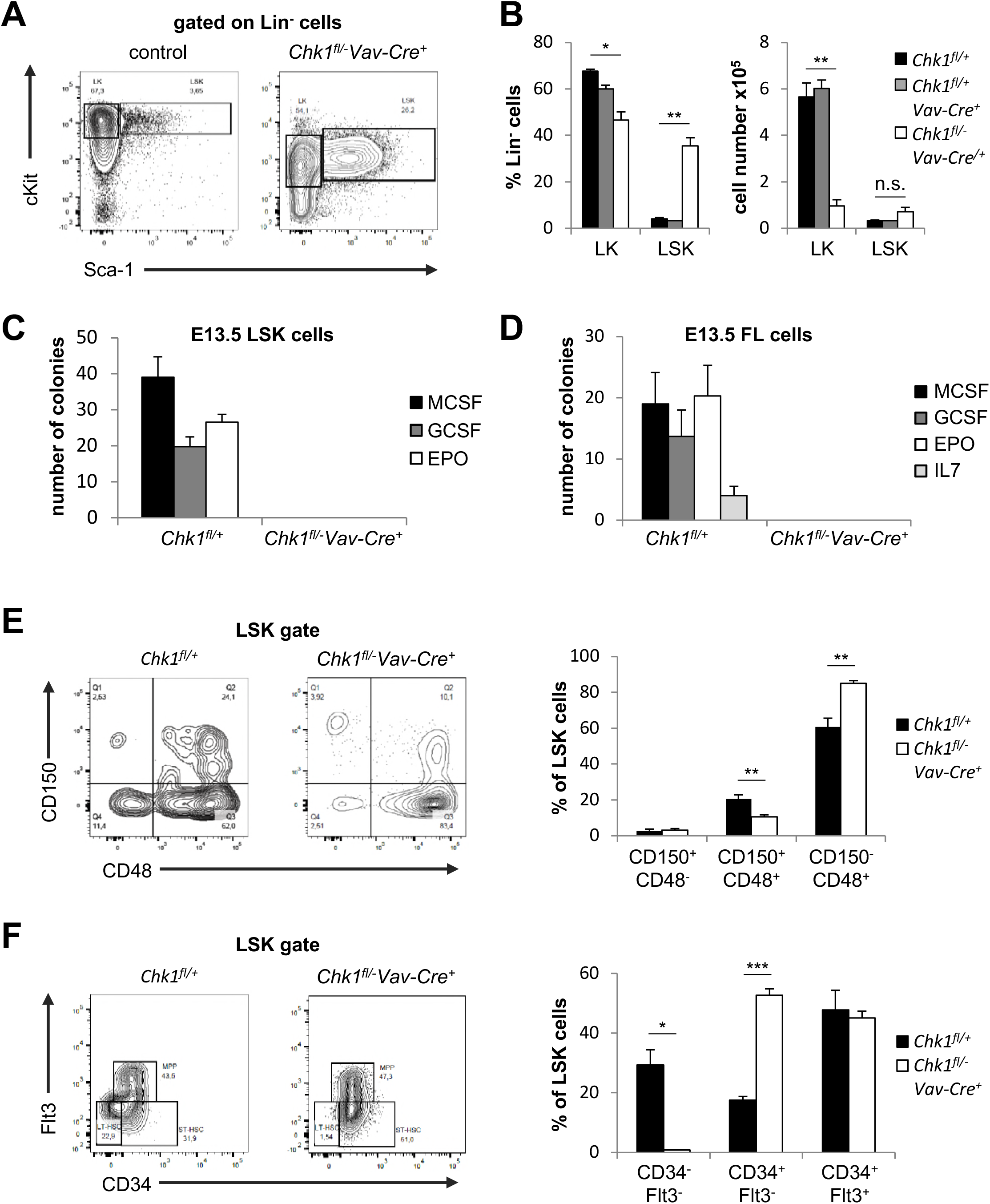
Loss of CHK1 causes signs of stem cell exhaustion in the fetal liver. **(A)** Representative dot-plots of the LSK cell phenotype observed in E13.5 fetal livers of *Chk1*^*fl/+*^ and *Chk1*^*fl/-*^ *Vav-Cre*^*+*^ embryos. **(B)** Relative distribution and absolute numbers of LK and LSK cells found in the fetal liver on E13.5. Bars represent means ± S.E.M. N=4 for *Chk1*^*fl/+*^, N=3 for *Chk1*^*fl/+*^ *Vav-Cre*^*+*^ and N=4 for *Chk1*^*fl/-*^ *Vav-Cre*^*+*^. **(C)** Colony formation potential of FACS-sorted LSK cells in methyl-cellulose assays. Bars represent means ± S.E.M. N=4 for *Chk1*^*fl/+*^ and N=5 for *Chk1*^*fl/-*^ *Vav-Cre*^*+*^. **(D)** Colony formation potential of fetal liver cells in methyl-cellulose assays. Bars represent means ± S.E.M. N=3 for *Chk1*^*fl/+*^ and N=6 for *Chk1*^*fl/-*^ *Vav-Cre*^*+*^. **(E)** Flow cytometric analysis of LSK cells stained with antibodies specific for CD150 and CD48 (N=6 for *Chk1*^*fl/+*^ and N=4 for *Chk1*^*fl/-*^ *Vav-Cre*^*+*^) or **(F)** CD34 and Flt3 (N=4 for *Chk1*^*fl/+*^ and N=4 for *Chk1*^*fl/-*^ *Vav-Cre*^*+*^), to discriminate LT- and ST-HSC from MPP within the LSK subset. Bars represent means ± S.E.M. Asterisks indicate significant differences: *p < 0.05, **p < 0.01, ***p < 0.001 using unpaired Student’s t-test.

To corroborate this phenotype further, we also performed colony formation assays in methyl-cellulose. Neither sorted LSK^low^ cells nor total FL cells isolated from *Chk1*^*fl/-*^*Vav-Cre* mice showed colony forming potential (Figure 5C,D). A more detailed analysis of the LSK cell pool indicated comparable percentages of CD150^+^CD48^-^ HSC and MPP1 cells, a decrease in CD150^+^CD48^+^ MPP2 paralleled by an increase in CD150^-^CD48^+^ MPP3/4 cells (Figure 5E). Using alternative markers to better define the HSC population, we noted signs of stem cell exhaustion, suggested by the lack of CD34^-^Flt3^-^ LT-HSC accompanied by a relative increase in CD34^+^Flt3^-^ ST-HSC. MPPs, characterized as CD34^+^Flt3^+^, where found unchanged using this maker combination (Figure 5F).

Together, this indicates that failure in blood cell development and embryonic lethality in *Chk1*^*fl/-*^ *Vav-Cre* mice is potentially due to exhaustion of LT-HSC that actively cycle to colonize the fetal liver after migrating in from the AGM region (6, 11). Stem cell exhaustion may be driven by the loss of ST-HSC, MPPs or their immediate progeny that, based on our in vitro findings above, may undergo apoptosis in the absence of CHK1.

### CHK1-loss causes compensatory proliferation and DNA-damage leading to stem cell exhaustion

Chemical inhibition of CHK1 induces DNA damage in cultured MPP-like cells and activates a p53-independent but mitochondrial cell death response (Figure 1). To explore if this is also responsible for the phenotypes seen *in vivo*, we analyzed FL cryo-sections by TUNEL-staining and FACS-sorted LSK cells and assessed signs of DNA-damage by intracellular staining for γH2AX. Consistent with our findings *in vitro*, we found a strong increase in TUNEL-positive cells in tissue sections and LSK as well as LK cells isolated from the fetal liver of *Chk1*^*f/-*^ *Vav-Cre* mice showed a strong increase in γH2AX phosphorylation (Figure 6A,B; Figure S2). Moreover, Ki67 staining suggested stem cell exhaustion (loss of G0 cells), due to increased proliferation of LSK and LK cells, forced to cycle potentially in an attempt to compensate for the loss of erythroid cells or their progenitors. Consistently, LSK as well as LK cells lacking CHK1 were devoid of Ki67 negative cells, indicating increased proliferation rates and simultaneously showed increased percentages of sub-G1 cells. Interestingly, the percentage of cells in S or G2/M-phase was unaltered (Figure 6C,D). Together, this suggests that these cells accumulate DNA damage while cycling in the absence of CHK1, leading to apoptosis, which could explain the loss of LK cells seen. The increased death of LSK cells appears to trigger increased proliferation in order to maintain absolute number and to generate more LK cells, but eventually leads to exhaustion (Figure 5).

**Figure 6:**
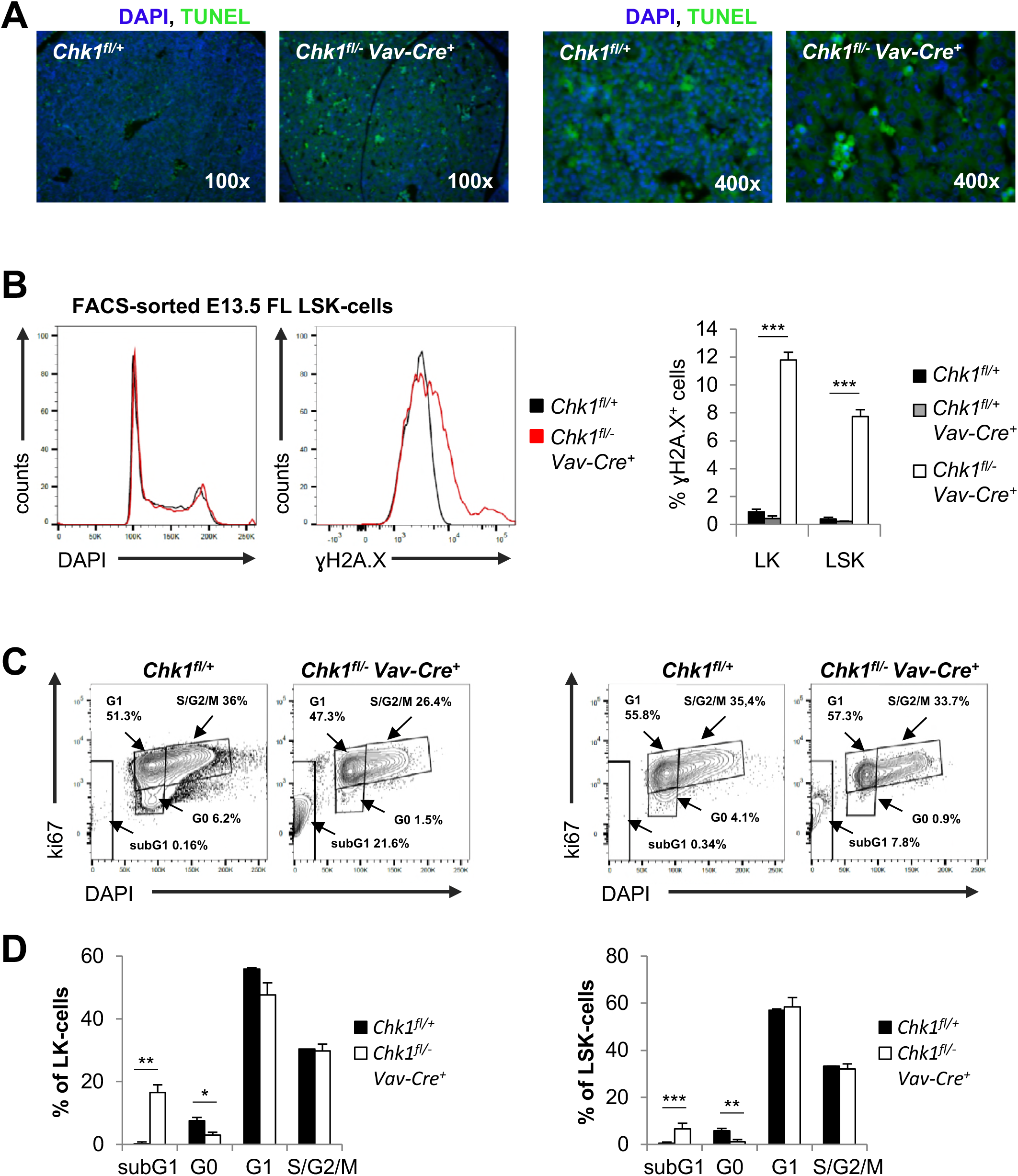
CHK1-deficient HSPCs accumulate DNA-damage during compensatory proliferation triggering cell death. **(A)** Terminal deoxynucleotidyl transferase dUTP nick end labeling (TUNEL) staining of fetal liver cryosections at E13.5. **(B)** LK- and LSK-cells of the indicated genotypes were sorted from E13.5 fetal livers and immediately fixed in 70% EtOH. Fixed cell suspensions were stained for γH2A.X and DAPI intracellularly. Shown here are representative histogram overlays and quantification of multiple analyses using N=4 animals per genotype. Total E13.5 fetal liver cell suspension were processed for intracellular Ki67 staining in combination with cell surface antibody staining to discriminate LK-from LSK-cells. **(C)** Representative dot-plots of Ki67 and DAPI staining in LK and LSK cells, quantified in **(D)** Bars represent means ± S.E.M. N=4 for *Chk1*^*fl/+*^ and N=5 for *Chk1*^*fl/-*^ *Vav-Cre*^*+*^. Asterisks indicate significant differences: *p < 0.05, **p < 0.01, ***p < 0.001 using unpaired Student’s t-test.

To assess the contribution of cell death to the embryonic lethal phenotype, we overexpressed BCL2 in fetal liver stem cells by intercrossing *Chk1*^*f/-*^ *Vav-Cre* with *Vav-BCL2* transgenic mice or by co-deleting *p53*. Consistent with a failure of *p53* loss to restore development of *Chk1*-deficient embryos (33, 34), we also failed to detect restoration of hematopoiesis in *Chk1*^*f/-*^ *Vav-Cre p53*^*-/-*^ *embryos*. Moreover, we also failed to observe a rescue of hematopoiesis upon BCL2 overexpression (Figure 7A,B), indicating that even if cell death was blocked, these cells may be unable to expand, as seen *in vitro* upon CHK1i treatment (Figure 1).

**Figure 7:**
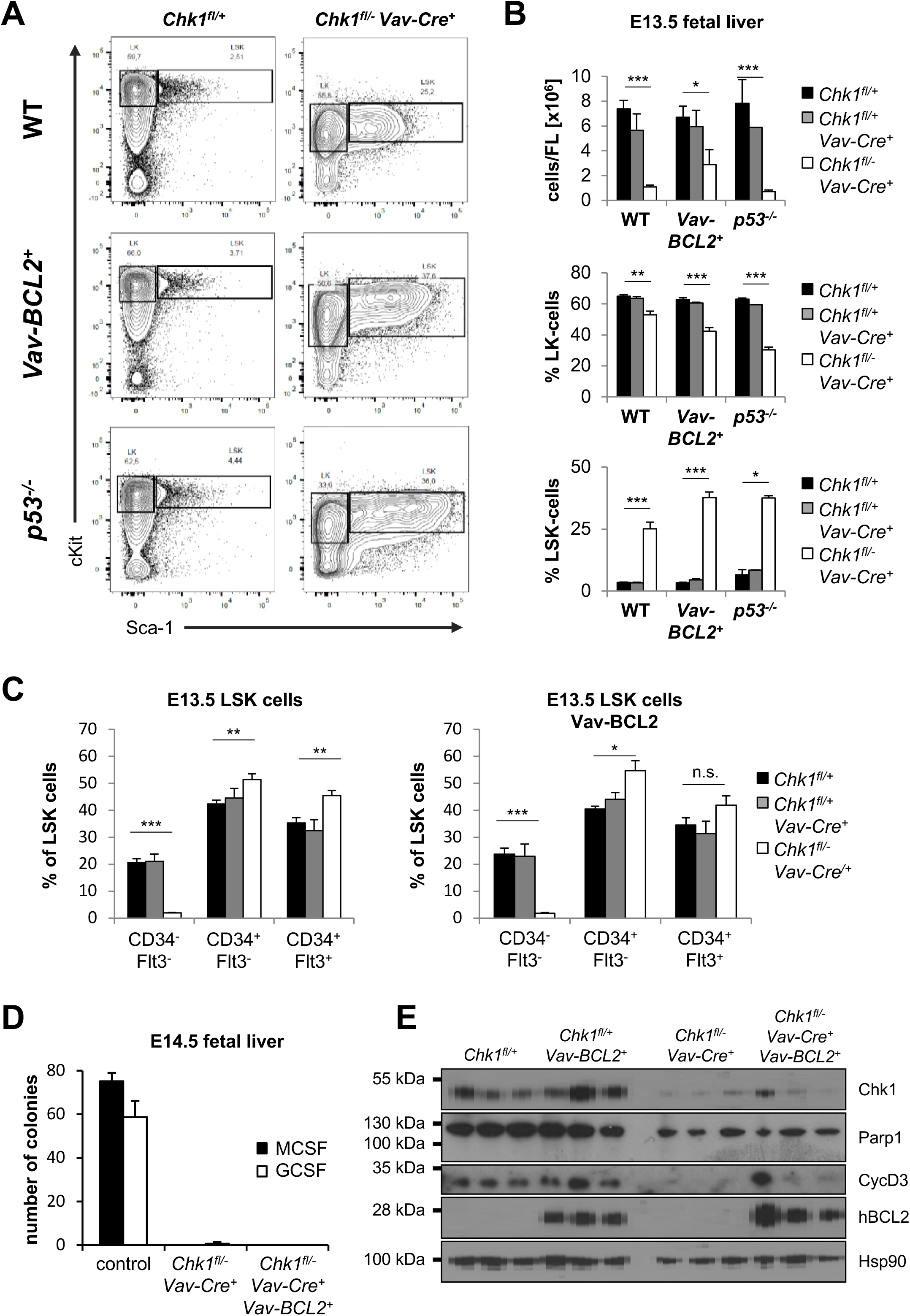
Apoptosis inhibition cannot restore hematopoiesis in the absence of CHK1. **(A)**Representative dot-plots and **(B)** quantification of E13.5 FL-derived LSK cell phenotypes observed in embryos of the indicated genotypes. N=21 for *Chk1*^*fl/+*^, N=5 for *Chk1*^*fl/+*^ *Vav-Cre*^*+*^, N=5 for *Chk1*^*fl/-*^ *Vav-Cre*^*+*^, N=16 for *Chk1*^*fl/+*^ *Vav-BCL2*^*+*^, N=7 for *Chk1*^*fl/+*^ *Vav-Cre*^*+*^ *Vav-BCL2*^*+*^, N=5 for *Chk1*^*fl/-*^ *Vav-Cre*^*+*^ *Vav-BCL2*^*+*^, N=2 for *p53*^*-/-*^ *Chk1*^*fl/+*^, N=1 for *p53*^*-/-*^ *Chk1*^*fl/+*^ *Vav-Cre*^*+*^, and N=4 for *p53*^*-/-*^ *Chk1*^*fl/-*^ *Vav-Cre*^*+*^. **(C)** Flow cytometry-based analysis of LSK cells stained with antibodies specific for CD150 and CD48 or CD34 and Flt3, to discriminate LT- and ST-HSC from MPP within the LSK subset in fetal livers of embryos of the indicated genotypes. N=25 for *Chk1*^*fl/+*^, N=7 for *Chk1*^*fl/+*^ *Vav-Cre*^*+*^, N=7 for *Chk1*^*fl/-*^ *Vav-Cre*^*+*^, N=16 for *Chk1*^*fl/+*^ *Vav-BCL2*^*+*^, N=6 for *Chk1*^*fl/+*^ *Vav-Cre*^*+*^ *Vav-BCL2*^*+*^, N=5 for *Chk1*^*fl/-*^ *Vav-Cre*^*+*^ *Vav-BCL2*^*+*^. **(D)** Analysis of the colony formation potential of fetal liver cells of the indicated genotypes in methylcellulose. N=4 for controls (1 *Chk1*^*fl/+*^, 2 *Chk1*^*fl/+*^ *Vav-Cre*^*+*^, 1 *Vav-BCL2*^*+*^), N=3 for *Chk1*^*fl/-*^ *Vav-Cre*^*+*^ and N=3 for *Chk1*^*fl/-*^ *Vav-Cre*^*+*^ *Vav-BCL2*^*+*^. **(E)** western blot analysis of fetal liver cells of the indicated genotypes at E14.5 using the indicated antibodies. Bars represent means ± S.E.M. with the exception for *p53*^*-/-*^ *Chk1*^*fl/+*^, and *p53*^*-/-*^ *Chk1*^*fl/+*^ *Vav-Cre*^*+*^. Asterisks indicate significant differences: *p < 0.05, **p < 0.01, ***p < 0.001 using unpaired Student’s t-test.

As seen before, we again noted a drop in LSK CD34^-^Flt3^-^ LT-HSCs combined with an increase in the percentage of LSK CD34^+^Flt3^-^ ST-HSCs and LSK CD34^+^Flt3^+^ MPPs. BCL2 overexpression did only modestly change the percentage of these cells. Despite an observable trend, the increase in cell number seen did not reach statistical significance (Figure 7B,C). Regardless, fetal liver cells overexpressing BCL2 were again unable to give rise to colonies in metho-cult assays (Figure 7D). This suggests that HSPCs that may survive the loss of CHK1 in the presence of high BCL2 levels fail to expand but undergo cell cycle arrest. Consistently, western analysis performed on E14.5 fetal liver cells confirmed that expression levels of CHK1 were drastically reduced in knockout embryos, as was the expression of Cyclin D, suggesting that the residual FL cells present were not cycling. Notably, BCL2 overexpression did not affect the loss of Cyclin D, confirming that when cell death is blocked, these cells fail to thrive.

Taken together, our findings suggest that HSPCs depend on CHK1 to prevent the accumulation of DNA damage during cell cycle progression during fetal liver colonization and that in its absence these cells undergo apoptosis. Yet, when intrinsic apoptosis is impaired, cell cycle arrest prevents HSPC expansion as a second barrier to prevent subsequent mutational spread.

## DISCUSSION

HSC are responsible for the supply of blood cells throughout our lifetime. Therefore, it is essential to control their stemness in response to proliferative cues to avoid stem cell exhaustion driving hematopoietic failure and to prevent accumulation of DNA damage or mutations that might contribute to malignant disease with age. CHK1 is a key-regulator of DNA-replication fidelity and the G2/M transition upon DNA damage, critical in early embryogenesis and meanwhile a recognized target in cancer therapy (23, 50). Yet, the impact of CHK1 loss or inhibition on normal hematopoiesis remains largely unknown.

We could recently show that human Burkitt’s lymphoma and pre B ALL cell lines, similar to primary pre-B and mature B cells from mice, die an apoptotic cell death controlled by the BCL2-family when treated with CHK1 inhibitors (51). Similarly, MPP-like Hoxb8-FL cells undergo mitochondrial apoptosis when treated with CHK1i. This death is clearly BAX/BAK dependent and can be delayed by BCL2 overexpression. Remarkably, despite the induction of DNA-damage and subsequent p53 stabilization, these cells are not protected from death by loss of p53. This suggests that NOXA and PUMA, two recognized pro-apoptotic p53 target genes acting upstream of BAX/BAK (52) may only play a limited role in this apoptotic response. The precise mechanism of BAX/BAK activation in response to CHK1 inhibition remain to be investigated in detail.

Our *in vivo* results clearly demonstrate that CHK1 is an essential component of the regulatory network controlling HSPC development and expansion. Conditional deletion of CHK1 in dHSC using Vav-CRE mediated recombination causes failure of fetal hematopoiesis, associated with a reduction in fetal liver cell number, loss of cKit expression and impaired hematopoiesis, detectable as early as on embryonic day 12.5 of gestation in *Chk1*^*f/-*^ *Vav-Cre* embryos (FS & AV; personal observations). Fetal livers of these embryos present with a relative accumulation of ST-HSC and MPPs, at the cost of LT-HSC that are driven into compensatory proliferation cycles and accumulate high levels of DNA damage. The lack of the colony formation capacity of DNA-damage baring fetal liver resident LSK cells that are cKit^low^ accounts for the embryonic lethality seen in *Chk1*^*f/-*^ *Vav-Cre* mice.

In support of this notion, we could detect only ~6% cKit^+^ cells in *Chk1*^*f/-*^ *Vav-Cre* fetal livers at E13.5 compared to more than 60% in littermate controls. Although cKit cannot be used as a general marker for dHSC and their progeny, as it is present on a variety of other cell types, it can be used to discriminate pHSC as well as primitive erythrocytes from dHSC and definite erythrocytes, as the former lack cKit, as reviewed in (6). Furthermore, it was shown that only cKit^+^ cells, but not cKit^low^ or cKit^-^ cells from the E11.5 AGM-region or the fetal liver can reconstitute lethally irradiated mice confirming that definitive hematopoiesis relies on the SCF-receptor, cKit (53). Together this suggests that loss of CHK1 using *Vav-Cre,* activated as early as on day E11.5 (54), results in the loss of definite hematopoiesis in the fetal liver and that the blood cells we can detect in *Chk1*^*f/-*^*Vav-Cre* embryos are of primitive hematopoietic ancestry. Interestingly, LSK cells isolated from fetal liver did not massively die in culture when treated with the CHK1-inhibitor while total fetal liver cells did die in a dose-, time- and BCL2-dependent manner, suggesting that loss of CHK1 may dominantly affect the survival of MPPs or their CLP or CMP progenitors. This may be explained by the fact that all these cells have a higher proliferative index compared to LSK cells. An alternative explanation would be the lack of CHK1 expression in fetal liver resident LSK cells, rendering them insensitive to inhibitor. Indeed, mRNA expression analysis showed lowest levels of *Chk1* mRNA in resting bone marrow LSK cells but higher levels in fetal liver LSKs as well as mobilized LSKs or MPPs (43), as well as increased protein in expanding human CD34^+^ HSPCs (47). Moreover, mobilized LSK cells showed clear signs of replication stress-associated CHK1 activation and hallmarks of DNA damage (44), indicating the need for CHK1 to deal with replication stress in cycling HSPCs that would otherwise trigger apoptosis.

This notion is further supported by the fact that we observed a 7-fold decrease in total fetal liver cell number in the absence of CHK1 but normal absolute numbers of LSK cells. In the absence of CHK1-mediated cell cycle control, LT-HSC lose self-renewal potential and exhaust because they seem to be recruited into the pool of transient amplifying ST-HSC and MPP. These cells then arrest their cell cycle in the presence of accumulating DNA damage when unable to die. Indeed, we could observe the induction of cell cycle arrest and *p21* mRNA when we tested BCL2-overexpressing or BAX/BAK-deficient Hoxb8-FL cells in response to CHK1i that cannot undergo apoptosis. Together, this might explain why overexpression of BCL2 in fetal liver HSPCs fails to rescue hematopoiesis in the absence of CHK1, despite protecting HSPCs from cell death *in vitro*.

Similar to what we noted in MPP-like cells and fetal liver cultures, deletion of CHK1 in the small-intestine, using CYP1A1 promoter to drive CRE expression in *AhCre* mice, leads to p53-independent apoptosis due to massive DNA damage. Remarkably, the GI tract can be restored in these mice from stem cells that fail to delete *Chk1* (55), a phenomenon not seen in *Chk1*^*f/-*^ *Vav-Cre* embryos. In line with these observations, developing mammary epithelial cells and early thymocytes do undergo cell death when CHK1 is deleted using *WAP-Cre* or *Lck-Cre*, respectively (37, 56). Furthermore, deletion of CHK1 in B cell progenitors using *Mb1-Cre* blocks B-cell development at the pro-B cell stage in the bone marrow (51). Of note, neither B cell nor T cell development could be restored by BCL2 overexpression, a phenomenon that might be due to the concomitant induction of cell cycle arrest. This response was documented in cycling pre-B cells unable to undergo apoptosis upon CHK1i treatment (36). Collectively this suggests that deletion of CHK1 is lethal for most cycling cell types, with the notable exception of adult hepatocytes that seem to be unaffected by loss of CHK1, most likely as homeostatic hepatocyte proliferation is minimal (57, 58), as well as chicken DT40 B lymphoma cells (59).

Of note, depletion of other components of the cell cycle network controlling faithful chromosome segregation and safe passage through M phase give phenotypes similar to those caused by CHK1 deletion. It was shown that loss of a single allele of the spindle assembly checkpoint component, *Mad2l1*, results in growth deficits of immature hematopoietic progenitor cells (60). Furthermore, deletion of *survivin*, a member of the chromosomal passenger complex (CPC) in adult hematopoietic cells results in bone marrow aplasia and mortality of hematopoietic progenitor cells whereas *survivin*^*+/-*^ mice show problems in erythropoiesis (61). Survivin-deficient hematopoietic progenitor cells fail to form colonies of the erythroid and megakaryocyte lineage (62). Moreover, conditional Aurora A kinase deletion using *Mx1-Cre* in a bone marrow reconstitution approach caused a severe cell-autonomous death in nearly all hematopoietic lineages (63). Together, this suggests that manipulation of the cell cycle machinery allowing for premature entry in or exit from mitosis results in hematopoietic failure or triggers a severe growth disadvantage in hematopoietic cells.

As inhibitors of CHK1 and related cell cycle regulating proteins such as Aurora kinase A or MAD2 are currently under investigation in clinical trials to treat various malignancies (as reviewed in (50)) it is important to understand if their transient inhibition can be tolerated temporarily or causes myeloablation, as seen frequently in response to chemotherapy. Conditional deletion of CHK1 or MAD2 in adult mice are still missing to address their suitability as drug-targets. *Chk1*^*+/-*^ animals on C57BL/6 background, however, were phenotypically normal and did also not show signs of anemia within their first year of life (F.S. & A.V.; personal observations), opening an opportunity to target cancer cells that appear to be even more dependent on CHK1 function for survival (32, 64). This idea is supported by a recent study testing a new CHK1i for its efficacy to kill AML in mice that proved particularly effective when combined with AraC and G-CSF (65). Here, the authors noted no negative side effects on the HSC or MPP compartment, suggesting that transient inhibition, in contrast to chronic (genetic) deletion, may indeed be tolerated, opening a window of opportunity for drug-treatment. Nonetheless, our findings raise the possibility that CHK1i treatment may become problematic during pregnancy as well as other situations that call stem cells out of dormancy, such as bacterial or viral infection, as well as chemotherapy-based combination regimens.

## Methods

### Mice

Animal experiments were performed in accordance with Austrian legislation (BMWF: 66-011/0106-WF/3b/2015). The generation and genotyping of *Chk1*^*f/f*^, *p53*^*-/-*^ *Vav-BCL2 and Vav-iCre* mice have been described (48, 56, 66-68). All mice were maintained or backcrossed on a C57BL/6N genetic background.

### Cell culture

All murine cells were cultured in RPMI-complete medium: RPMI-1640 medium (Sigma-Aldrich, R0883), supplemented with 10% FCS (Gibco, 10270-106), 2 mM L-glutamine (Sigma, G7513), 100 U/ml penicillin and 100 μg/ml streptomycin (Sigma, P0781) 50 μM 2-mercaptoethanol (Sigma, M3148). Hoxb8-FL cells were cultured in RPMI-complete supplemented with 1 µM β-estradiol and 5% supernatant of FLT-3L expressing B16-melanoma cells. Primary murine bone marrow cells, fetal liver cells or FACS-sorted bone marrow or fetal liver derived LK- or LSK-cells were cultured in RPMI-complete media, supplemented with 1 mM sodium pyruvate (Thermo Fisher, 11360070), 1x non-essential amino acids (Thermo Fisher, 11140035), 5% supernatant of FLT-3L expressing B16 melanoma cells and 2% supernatant of SCF-producing CHO (Chinese hamster ovary) cells. Human cord-blood derived CD34^+^ HSPCs were cultured in StemPro(tm)-34 SFM 1X serum-free medium (Gibco(tm); 10639011) supplemented with 10% ES-FCS (Gibco^TM^; 16141061), and human SCF (100ng/ml, Immunotools, 11343325), FLT-3L (100ng/ml; Immunotools 11340035), TPO (50ng/ml Immunotools, 11344863) and IL-3 (20ng/ml, Immunotools 11340035) cytokines.

### Generation of Hoxb8-FL cells

Fresh bone marrow suspension cells from adult mice were cultivated (1/10^th^ of the cells flushed from 1 tibia and 1 femur) for 2 days in Opti-MEM^TM^ (Gibco, Cat. 31985070) supplemented with 10% FCS, 2 mM L-glutamine, 100 U/ml penicillin, 100 μg/ml streptomycin, 50 μM 2-mercaptoethanol, 10 ng/ml IL-3 (Lot #120948 C2013, PeproTech), 20 ng/ml IL-6 (Lot #090850 A3013, PeproTech) and 2% supernatant of SCF-producing WEHI-231 cells. 2×10^5^ cells were then transduced via 3x 30 minutes spin-infection at 37 °C (500g, vortex in between) with the Hoxb8-encoding retrovirus using 250 µl Opti-MEM^TM^, 1.25µl metafectamin.

### Reagents

PF-477736 (Selleckchem S2904), CHIR-124 (Selleckchem S2683), QVD (SML0063, Sigma), DMSO (D5879, Sigma), Tamoxifen (Sigma, T5648).

### Flow cytometry and cell sorting

Flow-cytometric analysis or cell sorting of single cell suspensions generated from bone marrow or fetal liver was performed on an LSR-Fortessa or a FACS-Aria-III, respectively (both BD) and analyzed using FlowJo® v10 software. Antibodies used: ebioscience lineage-depletion antibodies (B220-bio RA3-6B2, Ter119-bio TER-119, CD3e-bio 145-2C11, CD11b-bio M1/70, Gr-1-bio RB6-8C5), NK1.1-bio (PK136, biolegend), Sca1-APC (D7, biolegend), Sca1-PE (D7, biolegend), cKit-PE/Cy7 (biolegend 2B8), cKit-FITC (biolegend 2B8), cKit-PerCP-Cy5.5 (biolegend 2B8), cKit-BV421 (biolegend 2B8), CD48-APC (HM48.1), CD150-PE/Cy7 (TC15-12), CD34-eFluor450 (RAM34, ebioscience), Flt3-PE (A2F10, ebioscience), CD45-PE (30-F11 biolegend), CD71-APC (R17217, ebioscience), B220-FITC (biolegend RA3-6B2), B220-APC/eF780 (ebioscience RA3-6B2), B220-PE (BD RA3-6B2), B220-PerCP-Cy5.5 (biolegend RA3-6B2), CD19-BV605 (biolegend 6D5), IgM-APC (biolegend RMM-1), IgM-FITC (BD II/41), IgM-eF450 (ebioscience eb121-15F9), IgD-PerCP-Cy5.5 (biolegend 11-26c2a), TCRβ-BV605 (BD H57-597), TCRβ-FITC (ebioscience H57-597), CD4-eF450 (ebioscience GK1.5), CD8-alexa647 (BD 557682), Mac1-APC (ebioscience M1/70), CD25-PE (biolegend PC61), CD93-PE/Cy7 (ebioscience AA4.1), NK1.1-APC (biolegend PK136), Ter119-PerCP/Cy5.5 (TER-119) AnnexinV-FITC (biolegend Lot: B206041), AnnexinV-eF450 (ebioscience; Lot: E11738-1633).

### Nicoletti staining

Cells were fixed in 1ml 70% ethanol while vortexing and stored at −20 °C for a minimum of 60min. Prior antibody staining, cells were washed twice (800g, 5’) with 2ml PBS to remove ethanol. After RNase A (Sigma) digestion (100 mg/ml in PBS, 30 min at 37 °C) cells were stained with propidium iodide (40 µg/ml) and the percentages of subG1, G1, S and G2/M cells were monitored by flow cytometry.

### Intracellular staining for γH2AX

Cells were fixed in ethanol and stored at −20 °C. After two washes with PBS, cells were incubated for 15min in PBS+0.25% Triton X-100 (Sigma; order #,) on ice for permeabilization. Cells were incubated for 60’ with anti-Human/Mouse phospho-H2A.X S139 mAb PerCP-eFluor® 710, (clone CR55T33, ebioscience). Cells were washed in PBS 1%BSA. DAPI (200 ng/ml) was used for DNA content analysis.

### Intracellular Ki67 staining

10^6^ bone marrow cells were incubated for 30 min at 4 °C with the primary antibody-mix: Lineage-biotin antibodies for B220, CD3, Gr1, Mac1, NK1.1, Ter119, plus antibodies recognizing Sca1-APC or cKit-PE/Cy7 (all 1/100 in PBS+10%FCS, 50 µl/tube). Cells were washed with 2ml PBS, then cells were stained with the secondary antibody-mix: streptavidin-PerCP/Cy5.5 (1/100 in PBS+10%FCS, 50 µl/tube). Cells were washed twice with 2ml PBS. Cells were fixed and permeabilized using the reagent from the APC-BrdU Flow-Kit (BD, Vienna, Austria), according to manufacturer’s recommendation: Cytofix/Cytoperm, 10’ on ice, 100µl/tube, Perm/Wash, 1ml/tube (400g, 5’), Cytofix/Cytoperm-PLUS, 10’ on ice, 100µl/tube, Perm/Wash, 1ml/tube (400g, 5’), Cytofix/Cytoperm, 5’ room temperature, Perm/Wash, 1ml/tube (400g, 5’). Then, cells were incubated for 30’ on ice with Ki67-alexa488 (16A8, biolegend), 1/100 in Perm/Wash buffer, 50µl/tube. Cells were washed twice with Perm/Wash buffer, 1ml/tube (400g, 5’), 300µl DAPI (final concentration 200 ng/ml) diluted in PBS+10%FCS was added and cells were analyzed on a flow cytometer.

### Viability assay

Cells were co-stained with Annexin-V-FITC, Annexin-V-Alexa647 (provider/#) or Annexin-V-Pacific blue (provider/#) 1:1000 in Annexin-V binding buffer; BD) and 7-AAD (1 µg/ml, Sigma). Cell death was analyzed by subsequent flow cytometric analysis.

### Isolation of CD34^+^ cells

Human cord blood was collected immediately following cesarean births after informed consent of the parents and approval of ethical committee of University Hospital Freiburg, Germany. After Ficoll density gradient-based separation of mononuclear cells, CD34^+^ cells were isolated with MACS technology (Miltenyl Biotec). The purity of CD34^+^ cells was generally greater than 90% as determined via FACS. The purified cells were then frozen in CryoStor CS 10 (Merck, C2874) and stored in liquid nitrogen for later use. After thawing, cells were cultured at a density of 2.5×10^6^ cells/ml overnight StemPro(tm)-34 SFM 1X serum-free medium (Gibco(tm), #10639011 supplemented with 10% ES-FBS (Gibco(tm), # 16141061) and human SCF (100ng/ml, Immunotools, 11343325), FLT3-L (100ng/ml; Immunotools, 11343305), TPO (50ng/ml Immunotools, 11344863) and IL-3 (20ng/ml, Immunotools, 11340035). For colony forming assays, MethoCult(tm) SF H4436 medium (Stem Cell Technologies) was used. Cells were added with or without CHK1 inhibitors at a density of 10^3^ cells/ml/35 mm cell culture dish and incubated for 10 days. Afterwards, colony counts and total cell counts were conducted. Lentiviruses GFP co-expressing pLeGO-iG lentiviral vector, Vsv.g-envelope and Gag/Pol plasmids were utilized for the production of BCL-2 expressing viruses. As described (69), HEK293T cells were used for viral packaging. CD34^+^ cells were incubated with lentiviruses in the presence of serum free medium with cytokines for 48 hours (MOI=10 per day).

### Gene expression

RNA was isolated from snap-frozen FACS-sorted or in vitro cultivated Hoxb8-FL cells using the Qiagen RNeasy Mini Kit (74104) and the RNase-Free DNase Set (79254) according to the manufacturer’s instructions. For each sample, hundred nanograms of RNA were reverse-transcribed to cDNA using the iScript cDNA Synthesis Kit (1708890) according to the manufacturer’s instructions. For qRT-PCR with the StepOne Plus System (Applied Biosystems), we used the 2x SYBR Green qPCR Master Mix from Biotool (B21203). Expression levels of *p21* mRNA were normalized to the housekeeping gene *Hprt*. Primers used were as follows (5’–3’): *p21*_Fw AAT TGG AGT CAG GCG CAG AT, *p21*_Rv CAT GAG CGC ATC GCA ATC AC, *Hprt*_Fw GTC ATG CCG ACC CGC AGT C *Hprt*_Rv GTC CTT CCA TAA TAG TCC ATG AGG AAT AAA C.

### Immunoblotting

Cells were lysed in 50 mM Tris pH 8.0, 150 mM NaCl, 0.5% NP-40, 50 mM NaF, 1 mM Na_3_VO_4_, 1 mM PMSF, one tablet protease inhibitors (EDTA free, Roche) per 10 ml and 30 μg/ml DNaseI (Sigma-Aldrich) and analyzed by Western blot analysis. For detection of proteins by chemoluminescence (Advansta, K-12049-D50) a mouse anti-CHK1 (CS 2360, 2G1D5), rabbit anti-γH2A.X (CS 2577), rabbit anti-p53 (Santa Cruz sc-6243), rabbit anti-PARP1 (CS 9542), mouse anti-HSP90 (Santa Cruz sc-13119), mouse anti-CycD3 (BD 554195), mouse anti-hBCL2 (clone S100) or rabbit anti-actin (CS 4967) were used. Goat anti-rabbit Ig/HRP (Dako, P0448) or rabbit anti-mouse Ig/HRP (Dako, P0161) were used as secondary reagents.

### Histology

For fixation and embedding E13.5 embryos were transferred to 4% paraformaldehyde (PFA in PBS) for fixation. Embryos were washed with 1 mM TBS (50 mM Tris-Cl, pH 7.5 150 mM NaCl) 2x 15 min, 2x 30 min, 70% EtOH, 80% EtOH and 90% EtOH each for 30 min followed by 3x 60 min incubation with 100% EtOH for dehydration. To remove the ethanol, embryos were incubated in methyl benzoate for a minimum of 24h. Embryos were incubated for 2x 30 min with benzol, then with a 1:1 benzol-paraffin mixture at 60 °C, then in molten paraffin at 60 °C. After positioning the embryos paraffin dishes were placed on a cooling plate to enable a quick and homogenous hardening of the paraffin. Histology blocks were cut sagittal with a rotation microtome (Reichert.Jung, Leica 2040) and sections pressed on silanized glass slides.

### TUNEL assay

Slides were incubated 3x 10 min with Xylene, 3x 5 min with 96% EtOH, 5 min with 70%, 50%, 30% EtOH and PBS for rehydration. Slides were re-fixed with 4% PFA in PBS for 20 min, washed with PBS for 30 min and permeabilized using 0.1% Triton X-100, 0.1% sodium citrate in a.d. for 2 min on ice. Slides were dehydrated after another washing step with TBS using 5 min incubation with 50%, 70%, 96% EtOH and rinsed in chloroform, then dried for 1 hour at 37 °C in a moist chamber. The TUNEL reaction was carried out by incubating sections with 15 µl of 0.6 µM FITC-labeled 12-dUTP (Roche, Basel, Switzerland), 60 µM dATP, 1 mM CaCl_2_, and terminal deoxynucleotidyl transferase buffer (30 mM Tris, pH 7.2, 140 mM sodium cacodylate), and 25 units of terminal deoxynucleotidyl transferase (Roche, Basel, Switzerland) for 1 hour at 37°C and covered with a plastic coverslip. The reaction was stopped by the addition of 10 mM Tris/1 mM EDTA, pH 8.0, for 5 minutes. Sections were washed twice in TBS before adding DAPI (2 µg/ml, in TBS, 40 µl/slide) for 10 min. Slides were washed twice with TBS and mounted in Mowiol^®^ 40-88 (Sigma-Aldrich) (15 µl/slide).

### Statistical analysis

Statistical analysis was performed using unpaired Student’s t-test or analysis of variance (ANOVA) for multiple group comparisons using Prism GraphPad software.

## Acknowledgements

We are grateful to K. Rossi, C. Soratroi, I. Gaggl, M. Fischer, J. Vier and J. Blitz for excellent technical assistance or animal care. We also thank J. Adams, S. Lowe, S. Elledge and T. Mak for sharing mouse models. This work was supported by the FWF-funded Doctoral College “Molecular Cell Biology and Oncology” (W1101) and grant # I3271, “New insights into the BCL2 family” (FOR2036) and by the Deutsche Forschungsgemeinschaft (grant HA2128/19-1). F. Schuler was supported by a Doc-fellowship from the Austrian Academy of Science (ÖAW).

## Author contributions

F.S. performed experiments, analysed data, prepared manuscript and figures, S.A. performed experiments with CD34+ HSPCs, M.E. supervised experiments with CD34^+^ HSPCs, G.H. provided Hoxb8-FL cells and related expertise, C.M. performed histological analysis, A.V. designed research, analysed data, wrote paper, conceived study.

---

**Supplementary Figure S1:**
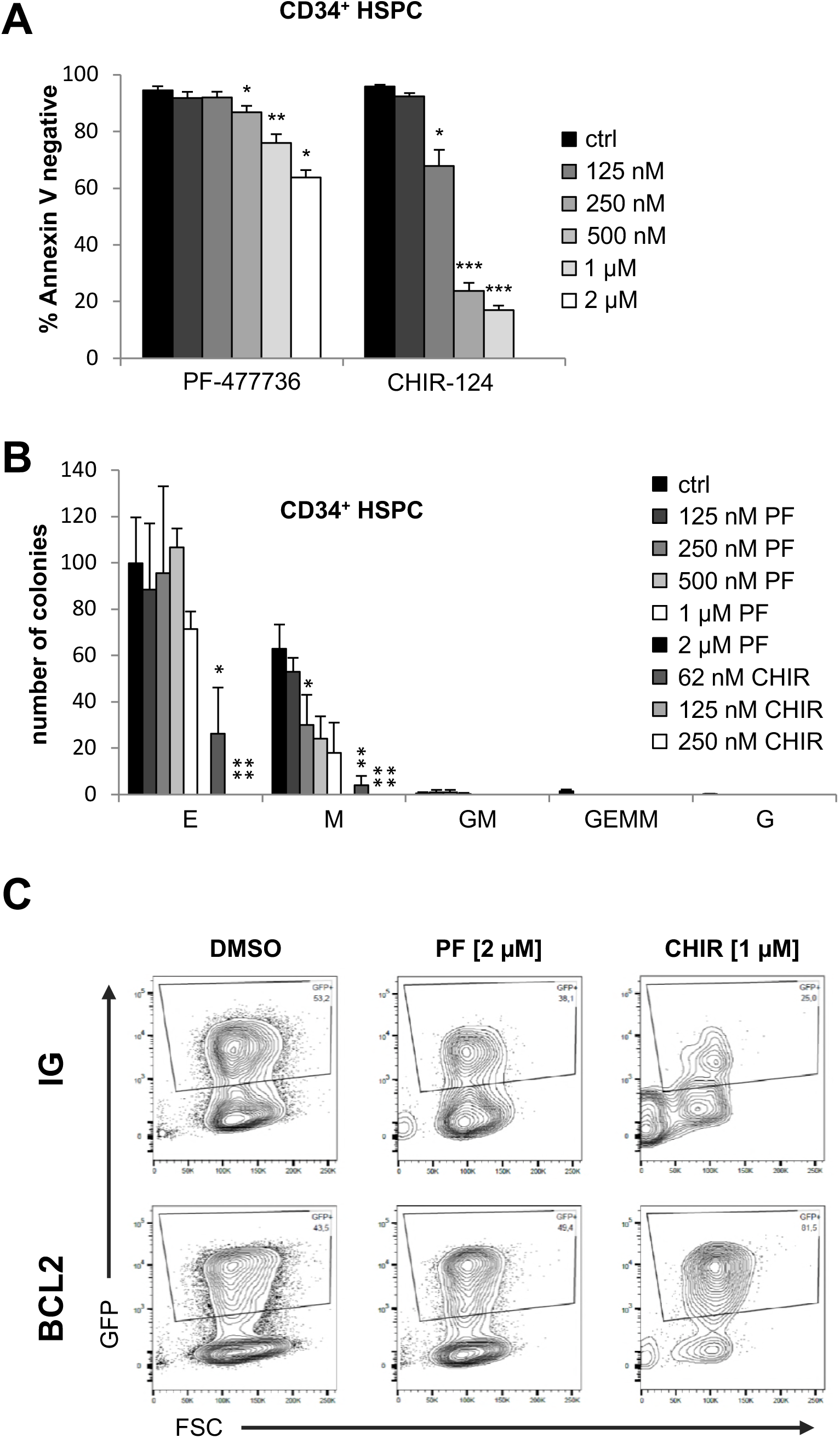
Human CD34^+^ HSPCs die an apoptotic cell death in response to CHK1-inhibition. **(A)** MACS-purified CD34^+^ human cord-blood derived HSPC were treated for 48h with graded doses of PF-477736 or CHIR-124. Cell death was assessed using AnnexinV/7-AAD staining and flow cytometry. N=4/genotype. **(B)** CD34^+^ HSPC were assessed for their colony formation capacity in metho-cult assays in the presence of graded doses of CHK1i. Colonies were counted 10 days post seeding of 1000 CD34^+^ cells. E: Erythroid, GM: granulocyte + monocyte, M: monocyte, G: granulocyte, GEMM: granulocyte, erythroid, monocyte/macrophage and megakaryocyte colonies. N=3-6 donors. Bars represent means ± S.E.M. Asterisks indicate significant differences: *p < 0.05, **p < 0.01, ***p < 0.001 using unpaired Student’s t-test. **(C)** Related to FIG3C: CD34^+^ cells were transduced with empty vector (IG) or a vector encoding BCL2, along with IRES-GFP followed by CHK1i application for 48 hours. Shown here are the percentages of GFP^+^ cells within living cells.

**Supplementary Figure S2:**
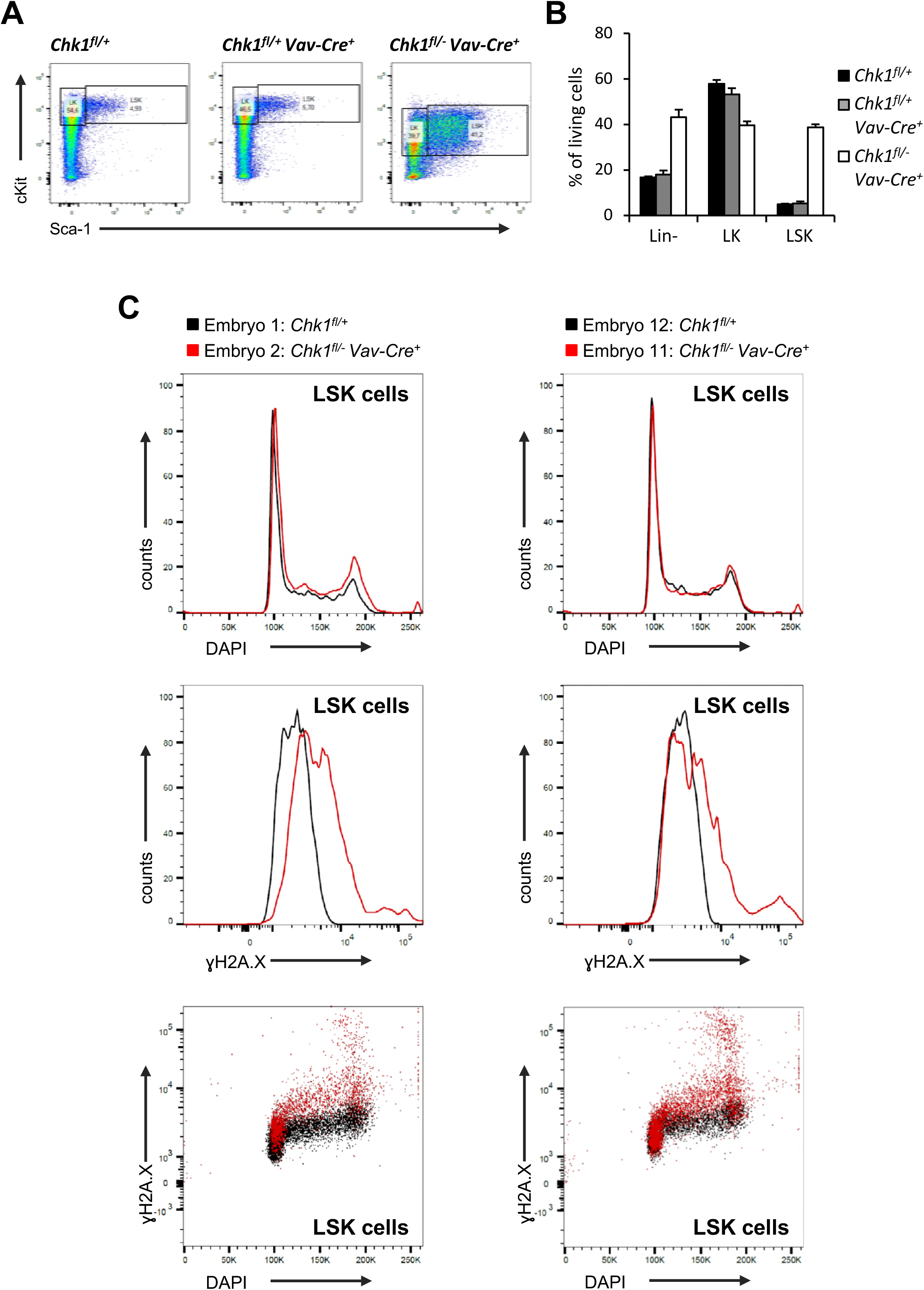
CHK1-deficient HSPCs accumulate DNA-damage. **(A)** Gating strategy for FACS-sorting LK and LSK cells for γH2A.X intracellular FACS and (B) quantification of Lin-, LK and LSK cells at E13.5. N=4/genotype. (C) Shown here are two additional examples of elevated γH2A.X-levels in *Chk1*^*fl/-*^ *Vav-Cre*^*+*^ embryos (red) at E13.5 as compared to *Chk1*^*fl/+*^ controls (black). ±S.E.M. *p < 0.05, **p < 0.01, ***p < 0.001 using unpaired Student’s t-test.

